# Reconstitution of immune cell interactions in free-standing membranes

**DOI:** 10.1101/311399

**Authors:** Edward Jenkins, Ana Mafalda Santos, James H. Felce, Deborah Hatherley, Michael L. Dustin, Simon J. Davis, Christian Eggeling, Erdinc Sezgin

## Abstract

The spatiotemporal regulation of signalling proteins at the contacts formed between immune cells and their targets determines how and when immune responses begin and end. It is important, therefore, to be able to elucidate molecular processes occurring at these interfaces. However, the detailed investigation of each component’s contribution to the formation and regulation of the contact is hampered by the complexity of cellular composition and architecture. Moreover, the transient nature of these interactions creates additional challenges, especially for using advanced imaging technology. One approach to circumventing these problems is to establish *in vitro* systems that faithfully mimic immune cell interactions, incorporating complexity that can be ‘dialled-in’ as needed. Here, we present an *in vitro* system making use of synthetic vesicles that mimic important aspects of immune cell surfaces. Using this system, we begin to investigate the spatial distribution of signalling molecules (receptors, kinases and phosphatases) and the intracellular rearrangements that accompany the initiation of signalling in T cells. The model system presented here is expected to be widely applicable.

**Summary Statement:** Immune cell-cell interactions are reconstituted in free-standing vesicles wherein spatiotemporal aspects of immune synapse formation can be investigated.

## Introduction

Dynamic cell-cell contacts govern the activation and effector functions of immune cells. Communication occurs through membrane protein interactions on opposing surfaces, whereby surface-presented antigens and ligands are recognised by key immune cell receptors, inducing intracellular signalling cascades and, eventually, the formation of an immunological synapse (IS), which comprises a spatiotemporally regulated supramolecular cluster of proteins at the interface between cells (Dustin and Choudhuri, 2016, Dustin and Baldari, 2017). Quantitative investigation of the key proteins and their molecular behaviour at the cellular contact is essential to gain insights into how immune cells integrate activating and inhibitory signals, allowing decisions about whether/when to respond (Dustin and Groves, 2012, Kamphorst et al., 2017). Studying these factors in physiological systems remains, however, challenging due to the intricacy and topographical complexity of cell-cell interactions. In addition, surface protein dynamics and organisation can be influenced by a variety of factors such as protein-protein or protein-lipid interactions, cortical actin cytoskeleton and glycocalyx, which makes it challenging to identify the exact role of each component (Chernomordik and Kozlov, 2003, Cho and Stahelin, 2005, Lemmon, 2008, Ritter et al., 2013). To this end, minimal *in vitro* systems with controllable complexity are useful for dissecting the determinants of the communication between cells.

The most basic systems to reconstitute immune cell interactions are planar substrates coated with immobile antibodies or purified biological ligands (Bunnell et al., 2001). Glass-supported lipid bilayers (SLBs) reconstituted with mobile proteins acting as surrogate antigen-presenting cell (APC) surfaces capture additional features of physiological T-cell/antigen presenting cell interfaces (Dustin et al., 2007). Advantages of SLBs include being able to control protein variety and density, and a simplified two-dimensional format allowing advanced optical imaging of the ‘cell-cell’ interface. As such, SLBs have been applied to study immune cell activation extensively (Bertolet and Liu, 2016, Dustin et al., 2007, Zheng et al., 2015, Lever et al., 2016). However, solid supports and SLBs suffer also from a number of disadvantages. First, the small hydration layer (1-2nm) between the bilayer and the underlying support is insufficient to completely de-couple the support’s influence on reconstituted proteins: the glass support restricts the diffusion of the molecules, mostly in an unpredictable manner, in the membrane plane thereby affecting the membrane dynamics significantly (Sezgin and Schwille, 2012, Przybylo et al., 2006).

Membrane fluidity in SLB-cell experiments has been shown to influence functional outcomes (Sanchez et al., 2015). Second, the solid glass support imposes rigidity on the lipid membrane. Although it varies, the stiffness of immune cell membranes is known to be several orders of magnitude lower than that of SLBs (0.1-1kPa vs. 1MPa; (Saitakis et al., 2017, Rosenbluth et al., 2006, Bufi et al., 2015), and substrate stiffness has been shown to influence B- and T-cell migration, synapse formation and signalling (Zeng et al., 2015, Shaheen et al., 2017, Natkanski et al., 2013, Judokusumo et al., 2012, Schaefer and Hordijk, 2015, Martinelli et al., 2014, Tabdanov et al., 2015). Lastly, the necessarily large area and planar nature of SLBs (*i.e*. centimetres) is a poor mimic of the topological constraints experienced by cells *in vivo* although this can be somewhat overcome by nanofabrication methods that partition bilayers (Choudhuri et al., 2014).

A simple alternative to SLBs is the giant unilamellar vesicle (GUV). These vesicular systems are not influenced by any surface (*i.e*., they are free-standing) and comprise a suitable mimic of cells with respect to their finite size (10-100 μm diameter), flexibility, deformability and free membrane fluctuations (Schmid et al., 2015, Fenz and Sengupta, 2012). Similar to SLBs, they can also be engineered to have various lipid compositions and to present membrane proteins, further emulating physiological membranes, as well as being amenable to light microscopy-based techniques. Very recently, GUVs have started to be employed to mimic T cells interacting with SLBs as the surrogate APC surface (Carbone et al., 2017). Minimal systems of this type (GUV-SLB) are likely to be especially informative with respect to understanding how spontaneous, topologically-driven processes shape the spatiotemporal properties of cell-cell contacts. At the next level, the analysis of live cells interacting with GUVs offers a way to dissect both passive and active processes driving the nascent immune response.

In this study, we utilised GUVs reconstituted with cell surface proteins as controllable, reductionist mimetics of APCs. Using this approach, we explored the interactions of T cells and mast cells with GUVs, directly observing the types of reorganisation of signalling proteins at the contacts that could explain the earliest stages of T cell and mast-cell activation.

## Results

### Preparation and characterisation of protein-loaded GUVs

SLBs are membrane bilayers sitting on top of a glass support with a thin water layer sandwiched between (Fig. 1A). Reconstitution of proteins on SLBs yields an effectively infinite flat surface that can be used as a mimic of an immune cell surface. Free-standing membranes, on the other hand, are freely floating vesicles that are not influenced by any surface (Fig. 1A) and have a finite size (typically tens of μm). Here, we explore the use of the free-standing giant unilamellar vesicles (GUVs) as a model system to mimic the immune cell surface.

**Figure 1.**
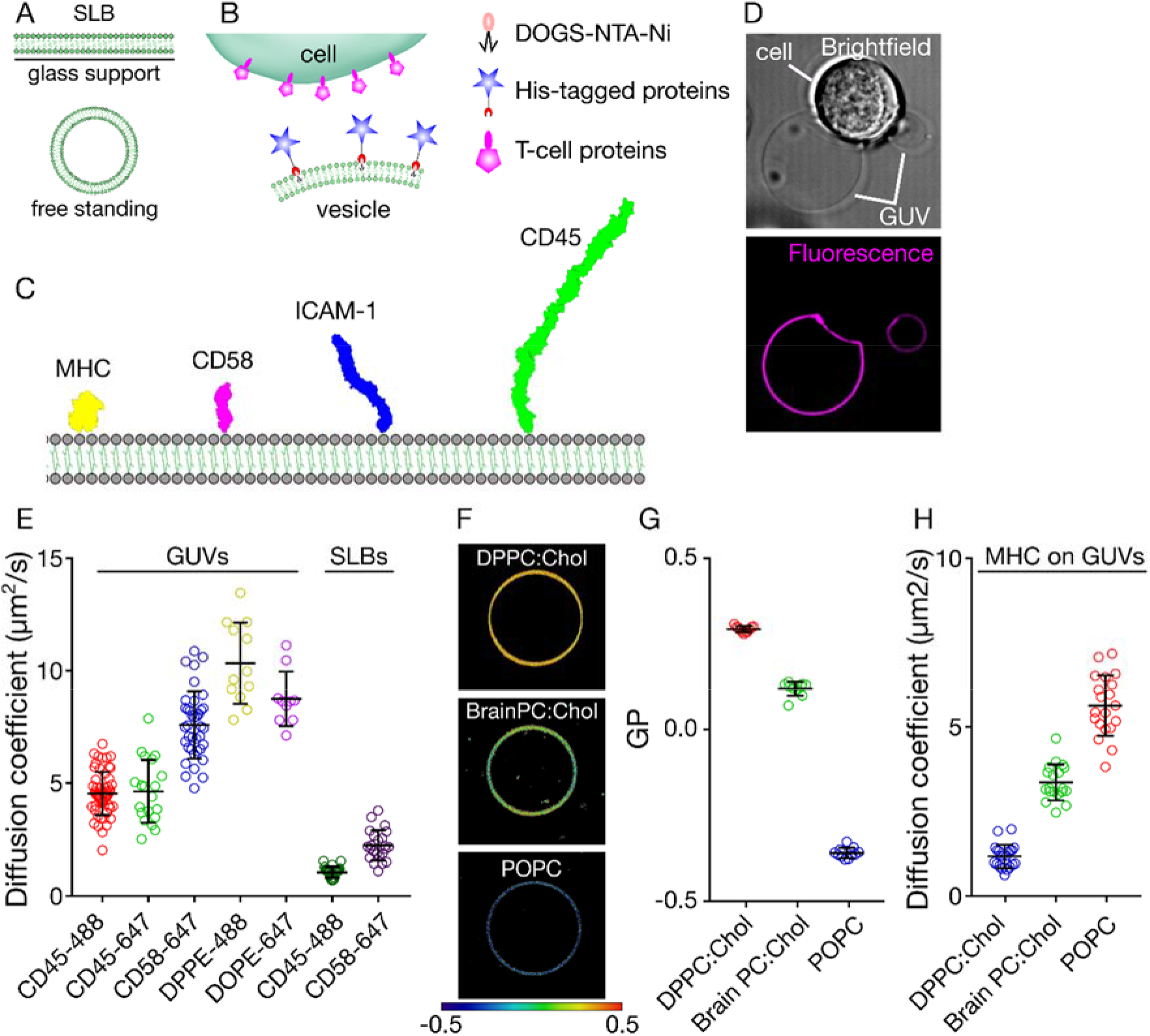
Scheme and the characterisation of the in vitro system. A) Schematic comparison of supported lipid bilayers with free standing vesicles. B) Scheme showing the in vitro cell/vesicle interaction. C) Molecules of interest for this study, drawn to scale based on structure determinations (Chang et al., 2016). D) Example bright field and fluorescence images of cell/vesicle contacts (image size 50 μm x 50 μm). E) Diffusion of fluorescently labelled lipids and proteins in GUVs and SLBs. F, G) Lipid packing of GUVs of varying composition revealed by a GP map and quantification of GP, respectively (image sizes are 40 μm x 40 μm). H) Diffusion of fluorescently labelled pMHC on GUVs composed of different lipids.

To attach immune cell surface proteins to the GUV surface, we prepared GUVs with a small fraction (4 mol%) of Ni-NTA functionalised lipids capable of directly binding to His-tagged surface proteins, each of which was expressed in a soluble form. To investigate protein interactions at the model cell-cell contact, we used His-tagged version of the proteins that are known to be involved in immune cell activation (Fig. 1B), *i.e*. pMHC, CD45, ICAM-1 and CD58 (Fig. 1C). With either pMHC or CD58 proteins presented on the GUVs, which are known to bind the T-cell receptor (TCR) and the small adhesion protein CD2 on the surface of T cells, respectively, we could successfully form cell/vesicle contacts (Fig. 1D).

As discussed above, reduced protein diffusion due to interactions with the glass support is a drawback of the SLB system. Given that interactions at the cell surface will be diffusion-limited once the membranes are in close proximity (Sanchez et al., 2015, Veya et al., 2015, Sezgin et al., 2017, Blouin et al., 2016), we were interested in characterising the diffusivity of the proteins on GUVs compared to SLBs. We measured the diffusion coefficients of proteins of interest using fluorescence correlation spectroscopy (FCS, see Materials and Methods for details). First, we tested whether the fluorescent tags have influence on protein mobility by comparing the diffusion coefficient of CD45 with different fluorophores (Alexa 488 and Alexa 647). Both of these molecules exhibited similar diffusion (Fig. 1E), confirming that there is no notable effect of the fluorescent tag on protein mobility. Next, we checked how the diffusion of proteins is affected by their size. Unsurprisingly, the smaller protein CD58 diffused much faster (almost as fast as lipids) than the larger protein, CD45 (Fig. 1E; for structure-based size comparisons see Fig. 1C). We next compared the diffusion of these proteins on GUVs versus SLBs. Strikingly, the diffusion of both proteins was significantly faster (~5 fold) in GUVs than in SLBs, presumably owing to influence of the glass support. Slower diffusion *i.e*. in more rigid membranes, is known to be achievable by changing the lipid composition of the vesicles (Machan and Hof, 2010). POPC is a phospholipid bearing saturated palmitic acid (16 carbon; 16:0), and mono-unsaturated oleic acid (18 carbon; 18:1) chains. Therefore, membranes composed of POPC are relatively fluid. Fluidity can be measured empirically using an index called generalised polarisation (GP) and polarity sensitive dyes (Sezgin et al., 2015b). A GP map shows lipid packing of membranes, with GP varying between −1 (representing maximally disordered membranes; dark blue) and +1 (representing maximally ordered membranes; dark red, see Materials and Methods for details of GP imaging). The fluidity of POPC GUVs, for example, is reflected by its blue colour in the GP map (Fig. 1F, G). Brain PC is a mixture of PC lipids (saturated and unsaturated), and cholesterol (Chol; which orders the membrane when present alongside unsaturated lipids), that yields a membrane of intermediate fluidity (Fig. 1F, G; yellow in the GP map). In contrast, DPPC carries two saturated palmitic acids, leading to the formation of a more rigid membrane. DPPC alone forms a gel phase, which is usually not found in biological systems. However, when cholesterol is present, DPPC forms a liquid phase with relatively low fluidity (Fig. 1F, G; red colour in GP map). To test the effects of these three membrane systems (POPC, Brain PC:Chol and DPPC:Chol) on protein diffusivity in the GUVs, we inserted the ligand of the TCR peptide; (p)MHC. We found that the pMHC complex diffused as fast as CD58 in POPC membrane (due to their similar sizes), nevertheless, its diffusion was slower in BrainPC:Chol and further reduced in DPPC:Chol membranes where it approached the diffusion rate of proteins in SLBs (Fig. 1H). This shows that, if required, more-saturated lipid mixtures can provide for more-constrained diffusion in free standing GUVs, slowing the diffusion to levels observed at native cell surfaces, as required. Hereafter, however, we use GUVs made only of POPC.

### Spatiotemporal reorganisation of cell surface proteins at cell/GUV contacts

The spatial organisation of signalling proteins at the cell-cell contact has been of great interest due to its likely role in the initiation of lymphocyte activation. Immune signalling events usually start with the phosphorylation of a receptor (such as the TCR or Fc receptors, FcRs) by intracellular kinases, such as Lck (Brownlie and Zamoyska, 2013). Yet, immune cells are also characterised by the very high expression of a large receptor-type protein tyrosine phosphatase, CD45, which, alongside other activities (most notably regulation of the kinases), is believed to antagonise the activities of kinases by dephosphorylating the receptors, keeping the cells in the resting state. Importantly, it has been shown that the TCR is at least as good a substrate for CD45 as Lck (Hui and Vale, 2014). Prior to cellular activation, the phosphatases and kinases are proposed to segregate from each other, insofar as the phosphatases are excluded from contacts of the antigen receptors with their ligands, but the kinases are not. It has been argued that the size-based exclusion of CD45 from contacts crucially breaks the balance between phosphatases and kinases in a way that favours the kinases (Davis and van der Merwe, 2006). Importantly, structural work on CD45 indicates that even the smallest form of CD45 is larger than the complex formed by the TCR and its pMHC ligands, suggesting a simple mechanism of steric exclusion (Chang et al., 2016). However, there have been other proposals for how the segregation of CD45 might occur *in vivo*, such as partitioning into lipid domains (Stone et al., 2017), charge effects (Su et al., 2016), or interactions of CD45 with active diffusional barriers created by integrins (Freeman et al., 2016).

The *in vitro* GUV-based system we present here is a useful tool for investigating the principles of protein spatial organisation at cell-cell contacts in three dimensions. We used the 1G4-TCR restricted T-cell line to study the formation of the contacts between cells and vesicles decorated with the His-tagged proteins shown in Fig. 1C, using the NTA-His coupling method depicted in Fig. 1B. These proteins were: (*i)* the pMHC recognised by the 1G4-TCR (i.e. a peptide derived from the tumour antigen NY-ESO (Chen et al., 2005)); (*ii)* CD58, which is the well-characterised ligand of the small adhesion protein, CD2; (*iii)* ICAM-1, which binds to LFA-1; and (*iv)* the phosphatase CD45, which is expressed by lymphocytes and also by antigen-presenting cells. The proteins were labelled fluorescently (colours are depicted in Fig. 1C; see Materials and Methods). We observed enrichment of pMHC and CD58 in the region of contact between the 1G4 T-cell and the GUV, as expected (Fig. 2A). The taller molecules, ICAM-1 and CD45, however, were excluded from the contact (Fig. 2A). Three dimensional images of the contacts in a large field of view further confirmed the segregation of CD45 and ICAM-1 from pMHC and CD58 (Fig. 2B), which could be quantified by fluorescence intensity line-profiling across the GUV (Fig. 2C; line shown in Fig. 2A). The ratio of the two peaks in the line profile, *i.e*. fluorescence at the contact side versus the non-contact side, was used to calculate the ratio of protein inside and outside the contact: “1” represents no preference, >1 represents enrichment inside the contact, and <1 represents exclusion from the contact. The inside/outside ratio was ~5-8 for CD58 and pMHC, and ~0.2-0.4 for CD45 and ICAM-1, confirming that CD45 and ICAM-1 are readily excluded from contacts enriched with CD58 and pMHC (Fig. 2D). To determine whether the pMHC/TCR interactions were needed for the exclusion of CD45, and if the T cell-expressed, versus the GUV-presented CD45 was excluded from the contact, we used GUVs presenting only CD2 and T cells expressing fluorescently labelled CD45 (i.e. the Fab fragment of Gap8.3 anti-CD45 antibody). Once again, we observed exclusion of CD45 from regions of the GUV surface where CD2 accumulated (Fig. 2E, F). This implies that a “close contact” between a T cell and a surface is sufficient to exclude CD45 at both sides of the contact. Since the GUV system is in thermodynamic equilibrium and lacks the actin cytoskeleton and lipid domains, the observed redistribution supports the idea that the proteins re-organise at cell-cell contacts largely according to size.

**Figure 2.**
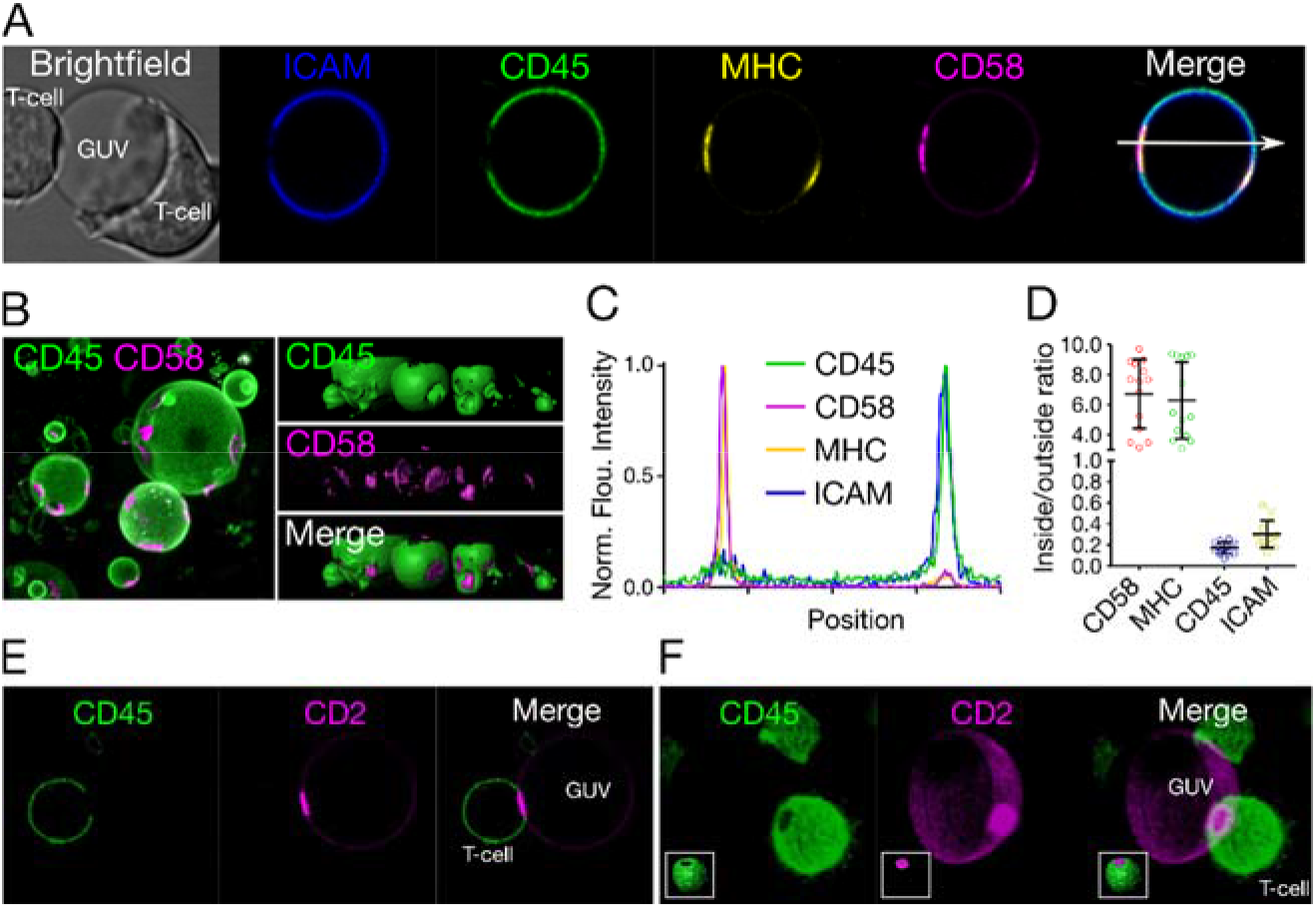
Protein reorganisation at cell/GUV contacts. A) Distribution of ICAM-1, CD45, pMHC and CD58 at cell/GUV contacts (image size 40 μm x 40 μm). B) 3D image (*left*, top view of raw image; and *right*, side view of surface image) of the contact formed between 1G4 T-cells and GUVs showing the abundance of contacts formed (image size 75 μm x 75 μm). C) Intensity line profile (line shown in panel A) of the fluorescence signal through the T cell contacting GUV. D) Quantification of the fluorescence signal inside and outside the contact zone (inside/outside ratio) for all four proteins. E) Relative distribution of CD45 (on the cell surface) and CD2 (decorating the GUV), showing that contact is sufficient for exclusion of CD45 phosphatase (labelled with anti-CD45 Gap8.3 Fab-Alexa488; image size 40 μm x 40 μm). F) Three-dimensional image of the contact shown in panel E (inset: side view surface image).

### Lymphocyte signalling induced by cell/GUV contact

In the course of lymphocyte activation, downstream-signalling tyrosine kinases are recruited to the triggered receptor. In T cells, this kinase is ZAP70, and in mast cells it is Syk (Brownlie and Zamoyska, 2013, Wernersson and Pejler, 2014). To test whether signalling could be induced in our GUV-based model system, we generated T-cell lines expressing Lck tagged with the fluorescent protein EGFP, and a form of ZAP70 that could be labelled with a HaloTag™. It is known that a pool of Lck is constitutively membrane-tethered *via* two palmitoylation and one myristoylation moieties. However, we were interested in determining whether Lck alters its distribution following receptor triggering and whether this could be induced by GUV-presented, TCR-triggering ligands. After incubating Lck-EGFP-expressing cells with GUVs presenting pMHC, we observed enrichment of Lck-EGFP in the region of contact (Fig. 3A; line scan shown in Fig. 3B). In the GUV-cell contact zone, we observed a clear recruitment of ZAP70, as well (Fig. 3C, D). ZAP70 recruitment is a robust indicator for cell activation, thus we can conclude that the free-standing *in vitro* system is able to successfully activate T cells.

**Figure 3.**
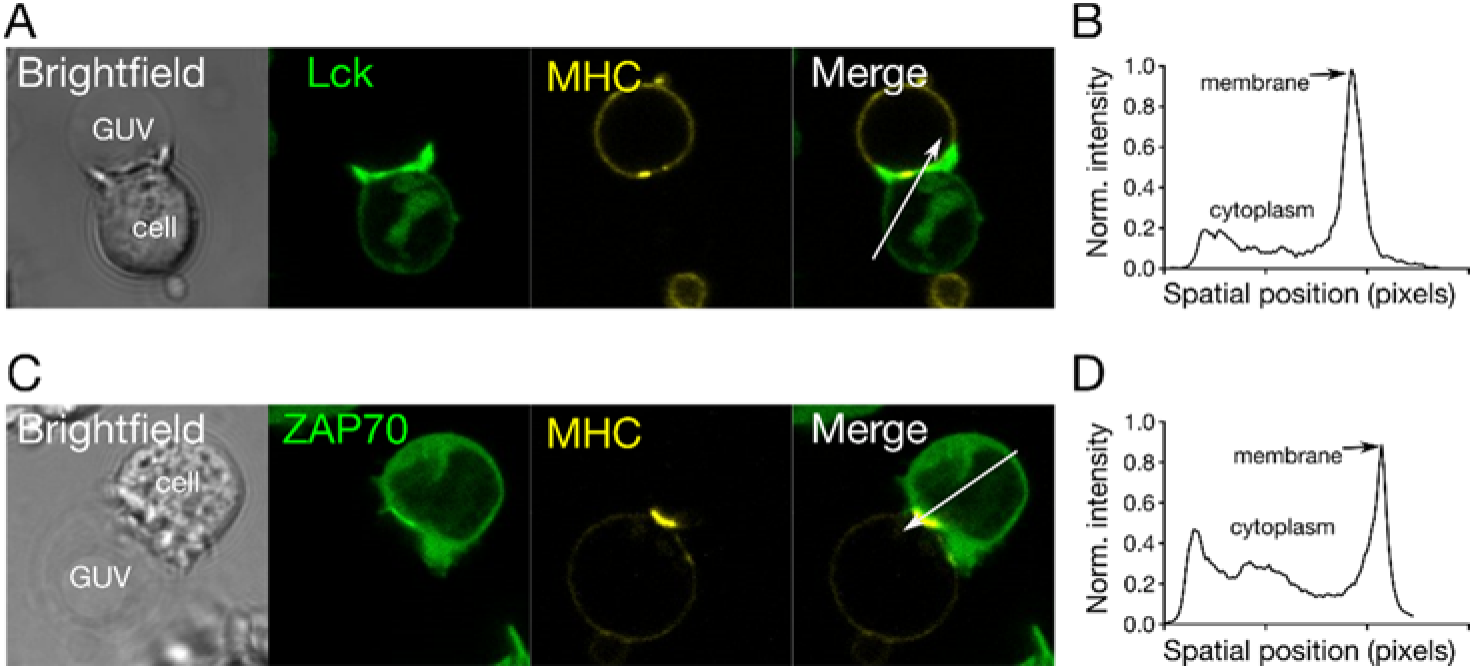
GUV-induced activation of T cells. A) Cellular localisation of Lck (labelled with EGFP, green) upon binding to GUVs decorated with pMHC. B) The intensity line profile of the Lck fluorescence signal through the contact shows enrichment of Lck at the contact zone. C) Cellular localisation of ZAP70 (labelled with HaloTag™) upon binding to vesicles carrying pMHC. D) The intensity line profile of the ZAP70-HaloTag fluorescence signal through the contact zone shows enrichment at the contact (image sizes 40 μm x 40 μm).

Finally, to investigate signalling in a non T-cell based system, demonstrating the versatility of the GUV/cell based model, we also tested whether we could observe FcεRI signalling in mast cells presented with GUVs decorated with receptor ligand mimics. We used rat basophilic leukaemia 2H3 (RBL-2H3; *i.e*. mast cells) expressing Syk kinases tagged with citrine fluorescent protein. To recreate model cell contacts, the GUVs were reconstituted with the Fc portions of rat IgE antibody (His-Fcε), which binds FcεRI on the mast-cell surface (Fig. 4A). FcεRI was itself labelled *via* a SNAP®-tag. When incubated with the GUVs, the RBL-2H3 cells similarly showed clear-cut recruitment of the Syk kinase to contacts where the receptor was also enriched (Fig. 4B, C).

**Figure 4.**
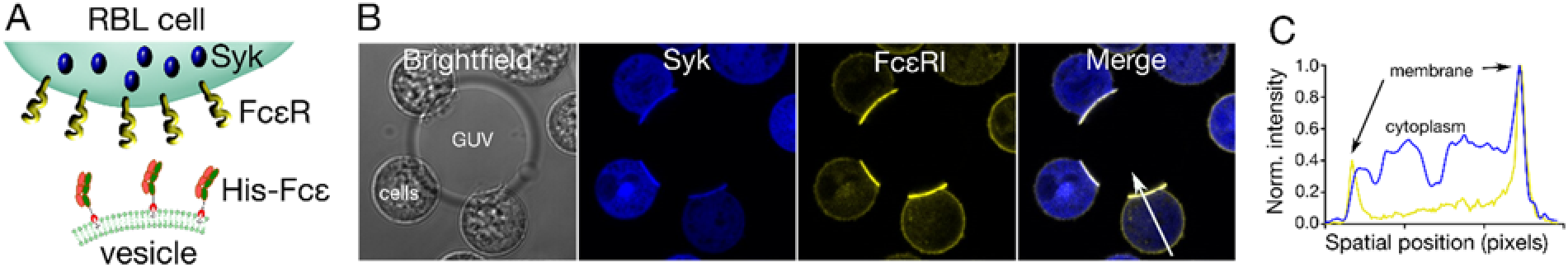
Mast-cell activation initiated by GUV/cell contact. A) Scheme showing the *in vitro* system used for studying RBL-2H3 (mast cell) signalling. Syk kinase (tagged with citrine) and FcεRI (labelled *via* a SNAP®-tag) were fluorescently labelled in the mast cells. GUVs were decorated with Fcε portions of the IgE antibody, which served as ligand for FcεRI. B) Example image of cellular localisation of Syk and FcεRI in RBL-2H3 cells upon contact with GUVs presenting Fcε ligand (image size 50 μm x 50 μm). C) Intensity line profile of the Syk and FceRI fluorescence signal through the contact (white arrow in panel B).

## Discussion

Reductionist *in vitro* reconstitution technologies are finding widespread use for analysing the biophysical basis of complex cellular processes (Jorgensen et al., 2017, Liu and Fletcher, 2009). GUVs have found applications in a broad range of fields, being used to probe many aspects of cell biology (Kahya, 2010, Bhatia et al., 2018, Prevost et al., 2017, Dubavik et al., 2012, Sezgin et al., 2015a, Richmond et al., 2011). Here, we presented a free-standing GUV-based membrane system with the potential to yield important insights into the spatiotemporal basis of immune cellcell interactions and lymphocyte activation.

We first showed that the GUV format allows the essentially unhindered diffusion of small and large proteins in the bilayer, giving diffusion constants substantially larger than those we obtained for SLBs. We also confirmed that it was possible to ‘fine-tune’ mobility by varying the lipid composition of the GUVs. By inserting key cell surface proteins expressed by antigen-presenting cells into GUVs and observing the contacts formed between the GUVs and live T-cells, we readily observed the patterns of large-scale spatial reorganisation of key surface proteins previously seen using supported lipid bilayers (Dustin and Choudhuri, 2016, Dustin and Groves, 2012, Dustin et al., 2007, Varma et al., 2006, Chang et al., 2016). Specifically, we observed the co-enrichment of small adhesion proteins, such as CD58, and activating ligands, such as pMHC, and the exclusion of large molecules and inhibitory proteins including ICAM-1 and CD45, respectively, at GUV/T-cell contacts. Surprisingly, however, we did not observe the formation of separate dSMACs and pSMACs, distal and peripheral regions of the contact characterized by the accumulation of CD45 and ICAM-1, respectively (Johnson et al., 2000, Freiberg et al., 2002), perhaps because T-cells need a more rigid substrate to form the well-organized radial lamellipodium necessary for dSMAC generation. Nevertheless, this minimal system was sufficient to induce early T-cell activation, as seen by the recruitment of downstream signalling effectors to the contact. We made analogous findings for mast cells triggered with the Fc portion of IgE presented by GUVs, reinforcing the similarities among leukocytes at least with respect to the earliest stages of signalling.

Importantly, we were able to confirm that the reorganisation of surface proteins occurred independently of cytoskeletal effects or the influence of lipid organisation in the GUVs, suggesting that receptor-ligand binding energies and the size-dependent lateral segregation of proteins were responsible for the observed patterning of molecules (Schmid et al., 2016). Importantly, despite the widespread interest in the possibility that the TCR acts as a mechanosensor (Brazin et al., 2015, Das et al., 2016), we observed contact formation, molecular re-organisation and signalling, all in the “softest” GUVs we could produce, implying that only very low levels of force are needed to effect signalling at cell-cell contacts. Although the reorganisation of CD58 and pMHC could in principle have been dependent on active processes occurring on the T-cell side of the contact, the behaviour of CD45 attached to GUVs was not, since there are no known ligands for CD45.

GUV-based systems provide useful frameworks for studying the functional outcomes of cell-cell contact in the immune system, offering the following advantages. The first, obvious advantage is that the 3D topology of a contact can be studied. This is especially important given the prominent role now assigned to microvilli-based contacts among immune cells (Jung et al., 2016). A second advantage, is that, as referred to above, the complexity and membrane properties of GUVs are highly tunable through the modular insertion of membrane proteins and by varying lipid mixtures. The use of more complex lipid mixtures or reconstitution of the actin cytoskeleton may provide a means to modify the stiffness of the GUVs. Since substrate stiffness has become increasingly important in understanding immune cell activation (Saitakis et al., 2017, Beningo and Wang, 2002), the GUV-based approach can be adapted to study this aspect of cell-cell contact. GUVs could be used, for example, to measure the influence of membrane tension on functional outcomes of immune cells based on membrane deformation. The third advantage is the finite size of GUVs, which make them better mimics of cells than SLBs. In principle, the size of unilamellar vesicles could be tuned to mimic smaller structures, such as organelles, *e.g*. microvilli, or microorganisms, pollen grains and viruses. Finally, in principle, vesicular systems allow the membrane reconstitution of full length proteins, including their transmembrane domains and intracellular regions, unlike SLBs, which opens up the possibility of studying the immediate sequelae of receptor triggering.

It is important, finally, to acknowledge the disadvantages of the GUV-based model system for studying cell-cell contacts. Since the vesicles are free-standing, they are not immobile over long-enough periods to allow hour-long measurements. Also, certain imaging technologies cannot be applied to GUVs, such as atomic force microscopy (AFM). Despite these disadvantages, which surely are not insurmountable, free-standing GUVs offer a powerful tool for dissecting cell surface biology both within and outside the immune system.

## Materials and Methods

### Lipids and Proteins

POPC, Brain PC, DPPC, DOGS-Ni-NTA and cholesterol are obtained from Avanti Polar Lipids. cDNA encoding ECD fragments of CD45 (CD45RABC, residues 24-575, UniProtKB P08575), CD58 (residues 29–215, UniProtKB P19265), CD54 (ICAM1, residues 28-480, UniProtKB P05362) and CD2 (ratCD2, residues 23-219, UniProtKB P08921) were ligated into pHR downstream of sequence encoding cRPTPσSP, having a ¾SRAWRHPQFGG¾‘spacer-his’ tag on the C-terminus. Soluble protein expressed lentivirally in 293T cells was purified using metal-chelate and size-exclusion chromatography. Soluble pMHC (HLA-A) was produced as previously described (Altman et al., 1996). For the purpose of labelling CD45 a fragment antigen-binding (Fab) digested from the whole antibody clone Gap8.3 tagged with Alexa 488 was used.

### Preparation of SLBs and GUVs

SLBs were prepared using spin-coating method (Clausen et al., 2015). #1.5 glass coverslips were first cleaned with piranha solution (sulfuric acid:hydrogen peroxide 3:1) for 45 minutes. After washing the coverslips with distilled water, 1 mg/ml lipid mixture (phospholipid:nickelated lipid, 96:4 molar ratio) was spread on them and immediately after they were spun with 3000 rpm for 40 seconds. Dried lipid bilayer was hydrated with SLB buffer (150 mM NaCl, 10 mM HEPES, pH 7.4). After the formation, SLBs were incubated with 1 μg/ml His-tagged protein for 30 minutes. Then, it is washed 10 times by first adding and removing fresh buffer.

GUVs were prepared using an electro-formation method. 1 mg/ml lipid mixture (phospholipid:nickelated lipid, 96:4 molar ratio) was deposited on platinum wire and dried. It was then dipped into a Teflon-coated chamber filled with 300 mM sucrose. GUV formation was triggered by a 10 Hz AC field for 1 h, which was followed by 2 Hz for 30 minutes. After the formation, 100 μL of the GUV suspension was incubated with 1 μg/ml His-tagged protein for 30 minutes. To wash out unbound protein, the GUV mixture was gently mixed with 1 ml PBS and allowed to sediment for 30 min. The bottom 100 μl was transferred to a new tube containing 1 ml PBS. This process was repeated twice. GUVs were imaged in PBS as described in the previous sections.

### Cell lines

Jurkat derived T-cell lines and were cultured in sterile RPMI supplemented with 10%FCS, 2 mM L-Glutamine, 1 mM Sodium Pyruvate, 10 mM HEPES, and 1% Penicillin-Streptomycin-Neomycin solution. Cells were maintained at 37 °C and 5% CO2 during culturing. Plasmid transfection and lentivirus infections were used to generate cell lines expressing the constructs described above (Lck-GFP, and ZAP-70-Halo).

RBL-2H3 cells (ATCC CRL-2256) were cultured at 37 °C, 5% CO2 in minimum essential medium eagle (MEM; Sigma-Aldrich) supplemented with 10% FCS, 1 mM L-glutamine (Sigma-Aldrich). 24 h before imaging, cells were incubated in falcon tube overnight on an end-over-end rotator (6 rpm) at 37 °C. FCεRI was labelled with SNAP tag and Syk labelled with citrine.

Halo and SNAP labelling was done using Oregon green-Halo (NEB) and JF-646 (kind gift from Janelia Farm laboratories), respectively. Cells were incubated with 0.1 mM (final concentration) of the dyes at 37 °C for 30 minutes. Afterwards, they were spun down at 1500 rpm for 3 minutes. Afterwards, they were washed by re-suspending in pre-warmed fresh medium (with all the supplements) and spinning down again. Later, fresh medium was added and they were incubated for another 30 minutes to get rid of unbound dyes inside the cells.

For cellular CD45 labelling, approximately 10^6^ cells were incubated with anti-CD45 (Gap8.3) Alexa Fluor 488-labeled Fabs (degree of labelling ~ 2 moles of dye per moles of protein) diluted in 100 μl of Hepes buffer at a final concentration of 10 nM at 37 °C for 15 minutes.

### Fluorescence correlation spectroscopy (FCS)

FCS on the GUVs and SLBs were carried out using Zeiss 880 microscope, 40X water immersion objective (numerical aperture 1.2) as described before (Schneider et al., 2017). Briefly, before the measurement, the shape and the size of the focal spot was calibrated using Alexa 488 and Alexa 647 dyes in water in an 8-well glass bottom (#1.5) chamber. To measure the diffusion on the membrane, the SLBs formed on #1.5 glass or GUVs placed into an 8-well glass bottom (#1.5) chamber were used. The laser spot was focused on the membrane by maximising the fluorescence intensity. Then, three curves were obtained for each spot (five seconds each). The obtained curves were fit using the freely available FoCuS software (Waithe et al., 2015).

### Confocal and spectral generalized polarisation (GP) imaging

After GUVs are gently transferred to an 8-well Ibidi chamber filled with 250 μL PBS, 50 μL of cells suspended in Fluorobright medium (a low fluorescence version of standard DMEM medium) was added into the wells. The imaging is performed at 37 °C in PBS. The samples were imaged with a Zeiss LSM 780 or 880 confocal microscope. Pacific Blue were excited with 405 nm and emission was collected with band pass 420-480 nm. Alexa 488-labeled proteins were excited with 488 nm and emission collected between 505-550 nm. Alexa 555-labelled proteins were excited with 543 nm and emission is collected between 570-630 nm. Alexa 647-labeled molecules were excited with 633 nm and emission collected with LP 650 filter. Multi-track mode was used to eliminate the cross talk between channels. Images are later anayzed in FiJi/ImageJ. For 3D images, Imaris is also used.

Spectral imaging was performed on a Zeiss LSM 780 confocal microscope equipped with a 32-channel GaAsP detector array. Laser light at 405 nm was selected for fluorescence excitation of Laurdan. The lambda detection range was set between 415 and 691 nm for Laurdan. The images were saved in .lsm file format and then analyzed by using a freely available plug-in compatible with Fiji/ImageJ, as described in ref (Sezgin et al., 2015b).

## Acknowledgement

We would like to thank Wolfson Imaging Centre for providing imaging tools, Dominic Waithe for his help on image processing, Yuan (Oliver) Lui for his help on protein drawings and Jemma McBride for her help on recombinant proteins.

## Conflict of Interest statement

Authors declare no conflict of interest.

## Funding

E.S. is supported by EMBO long term (ALTF 636-2013), Marie Skłodowska-Curie Intra-European Fellowships (MEMBRANE DYNAMICS-627088) and Newton-Katip Celebi Institutional Links grant (352333122). This work is supported by the Wolfson Foundation (ref. 18272), the Medical Research Council (MRC, grant number MC_UU_12010/unit programmes G0902418 and MC_UU_12025), MRC/BBSRC/ESPRC (grant number MR/K01577X/1), the Wellcome Trust (grant refs. 104924/14/Z/14, 100262Z/12/Z and 098274/Z/12/Z), Deutsche Forschungsgemeinschaft (Research unit 1905), and internal University of Oxford funding (EPA Cephalosporin Fund and John Fell Fund). J.H.F is supported by a Wellcome Trust Sir Henry Wellcome Fellowship (107375/Z/15/Z).

## Data Availability

The data is available.

